# Morphology and Blood Metabolites Reflect Recent Spatial Differences Among Lake Winnipeg Walleye, *Sander vitreus*

**DOI:** 10.1101/2020.04.10.035832

**Authors:** Matt J. Thorstensen, Lilian M. Wiens, Jennifer D. Jeffrey, Geoffrey M. Klein, Ken M. Jeffries, Jason R. Treberg

## Abstract

The invasive rainbow smelt (*Osmerus mordax*) was an abundant food source for Lake Winnipeg walleye (*Sander vitreus*), especially in the north basin of the lake, until the smelt’s collapse in approximately 2013. We quantified changing length-at-age (≈ growth rates) and relative mass (≈ body condition) in Lake Winnipeg walleye caught for a gillnet index data set. Here, walleye showed smaller length-at-age, particularly in the north basin with young fish, over time. This approach to assessing growth suggests a constraint in the north basin fish, possibly a nutritional limitation between 2017 and 2018, that was not present in the south. We then analyzed a separate group of walleye (≥452 mm in fork length) sampled in 2017 as part of a large-scale tracking study, which had a similar slope in length-mass relationship to large walleye caught in that year for the gillnet index data. A panel of metabolites associated with amino acid metabolism and protein turnover was compared in whole blood. These metabolites revealed elevated essential amino acids and suggest protein degradation may be elevated in north basin walleye. Therefore, based on both growth estimates and metabolites associated with protein balance, we suggest there were spatially distinct separations affecting Lake Winnipeg walleye with decreased nutritional status of walleye in the north basin of Lake Winnipeg being of particular concern.

## Introduction

Walleye (*Sander vitreus*) are the largest component of the Lake Winnipeg fishery in Manitoba, the second-largest freshwater fishery in Canada (Fisheries and Oceans Canada, 2018). Sustainable management of these walleye is, therefore, of enormous importance to commercial fishing, recreational angling, and the Lake Winnipeg ecosystem. However, annual commercial yield from the Lake Winnipeg walleye fishery has been above maximum sustainable yield since 2002, while commercial harvests have also declined between 2014 and 2018 (Manitoba Sustainable Development, 2018). Relatively recent observations of dwarf walleye, primarily in the south basin, suggest a selective pressure against large individuals—a selective force possibly induced by fisheries (Moles et al., 2010; Sheppard et al., 2018). Taken together, these separate pieces of evidence indicate that the important Lake Winnipeg walleye fishery is faced with several issues that may affect its sustainability and suggest that this fishery may need conservation attention.

To address conservation issues such as those that the Lake Winnipeg walleye may face, resource managers require many pieces of information to forecast how actions affect the probability of distinct alternative futures (Gattuso et al., 2015; Dudgeon et al., 2006). Simultaneous exposure to multiple stressors, such as eutrophication and invasive species, may lead to cumulative detrimental impacts on a fishery (Schindler et al., 2001). One key alteration to the Lake Winnipeg ecosystem was the introduction and subsequent crash of non-native rainbow smelt (*Osmerus mordax*). The rainbow smelt were first been observed in Lake Winnipeg in late 1990 (Franzin et al., 1994), and were later found in the stomachs of 82.9% of walleye caught in the north basin, but only 9.3% of walleye caught in the south basin (in 2010 and 2011, see Sheppard et al., 2015). At present however, rainbow smelt have almost disappeared from Lake Winnipeg, and their disappearance coincides with walleye body condition (a measure of mass relative to length, or ‘fatness’ of the fish) declines across the lake (Caskenette et al., submitted in this issue; Enders et al., submitted in this issue; Manitoba Government, 2018). As walleye body condition decreases over time, implicating a reduction in available nutritional resources such as the rainbow smelt, while fishing effort remains constant or rises, the recent declines in Lake Winnipeg walleye abundance may be exacerbated (Manitoba Sustainable Development, 2018). Therefore, understanding how changing food availability may be affecting Lake Winnipeg walleye is a fundamental issue in the system and a useful piece of information for resource management.

However, beyond gross morphological measurements like body condition, there are few available tools for evaluating the nutritional status of wild fishes outside of destructive sampling and proximate composition analysis. The current study was undertaken, taking advantage of the patterns of declining fish condition, to determine if there may be basin-level differences in the nutritional status of walleye in Lake Winnipeg by examining body condition and growth rates over time. Basin-level differences seemed most likely because the lake is characterized by two large basins separated by a narrow channel (Figure 1). Our second goal was to identify potential biomarkers from non-lethal sampling to test for differential nutritional status or energetic demands in walleye from the different regions of the lake. With refinement and validation, such biomarkers could provide managers with tools that may be able to rapidly inform on physiologically relevant thresholds of nutritional constraints that would be useful in risk assessments (Connon et al., 2018).

**Figure 1.**
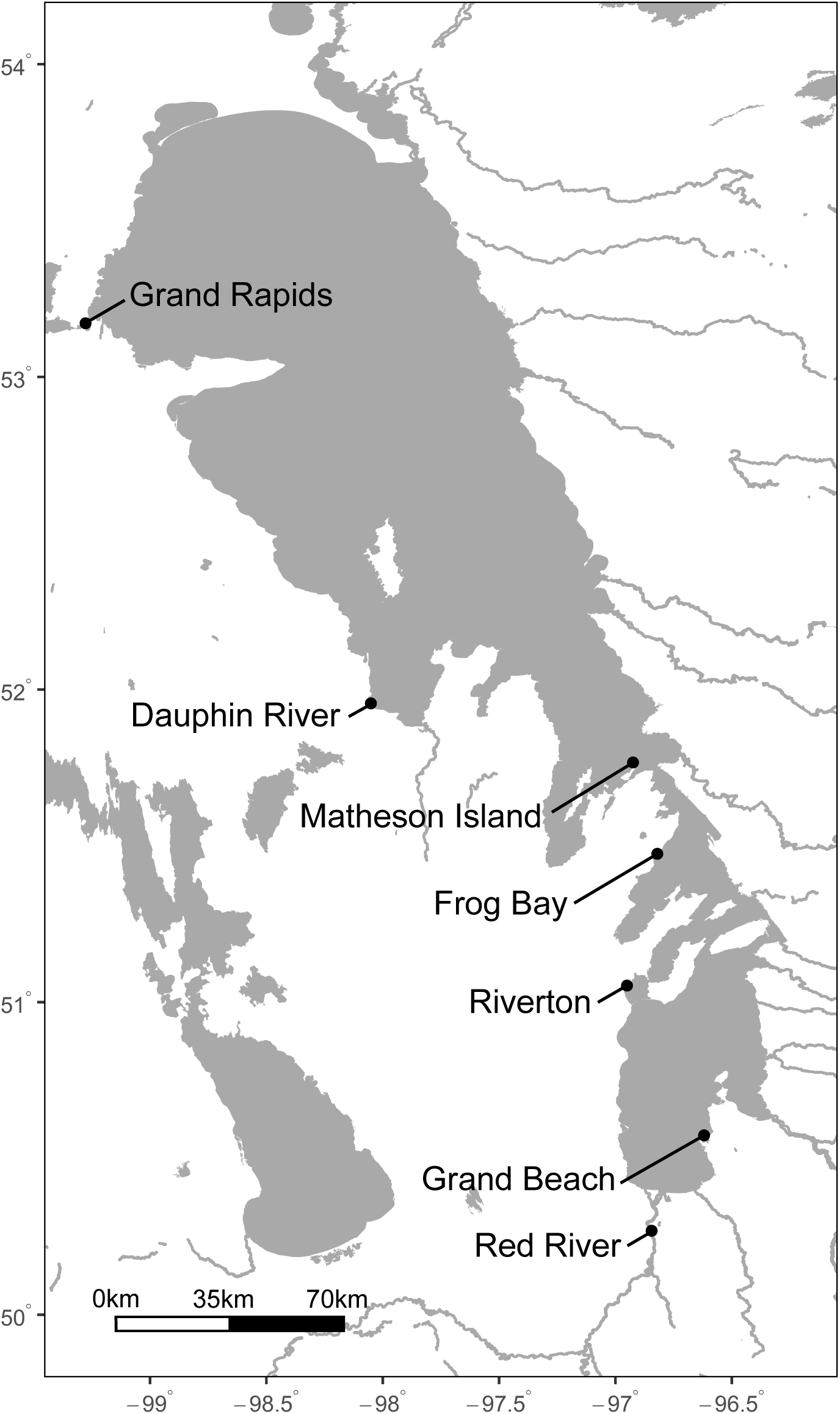
Map of Lake Winnipeg and the sites included in the present study. The Grand Rapids and Dauphin River represent the north basin, Matheson Island and Frog Bay the channel, and Riverton and Grand Beach the south basin, in the gillnet index data collected between 2009 and 2018 by the Government of Manitoba. For the blood metabolite data consisting of large walleye (*Sander vitreus*) caught by electrofishing in 2017, the Dauphin River represents the north basin, Matheson Island the channel, and the Red River the south basin.

### Nutritional biomarker strategy

We focused our study on protein metabolism because bulk protein growth is, with certain caveats, analogous to individual fish growth (see Carter and Houlihan, 2001 for review). Protein growth is a function of the balance between the rate of synthesis of new proteins from free amino acids and the rate of protein degradation, which releases free amino acids (Figure 2). Therefore, if an animal is synthesizing more protein than the rate of protein breakdown, there is net growth. Protein synthesis and degradation are both tightly regulated physiological processes, and as such the balance between both processes are intimately linked to growth and energy balance. We sought a biomarker strategy that could differentiate between amino acid breakdown for oxidation as an energy source (as opposed to re-using amino acids for protein synthesis), and the extent of endogenous protein breakdown in walleye across Lake Winnipeg (Figure 1).

**Figure 2.**
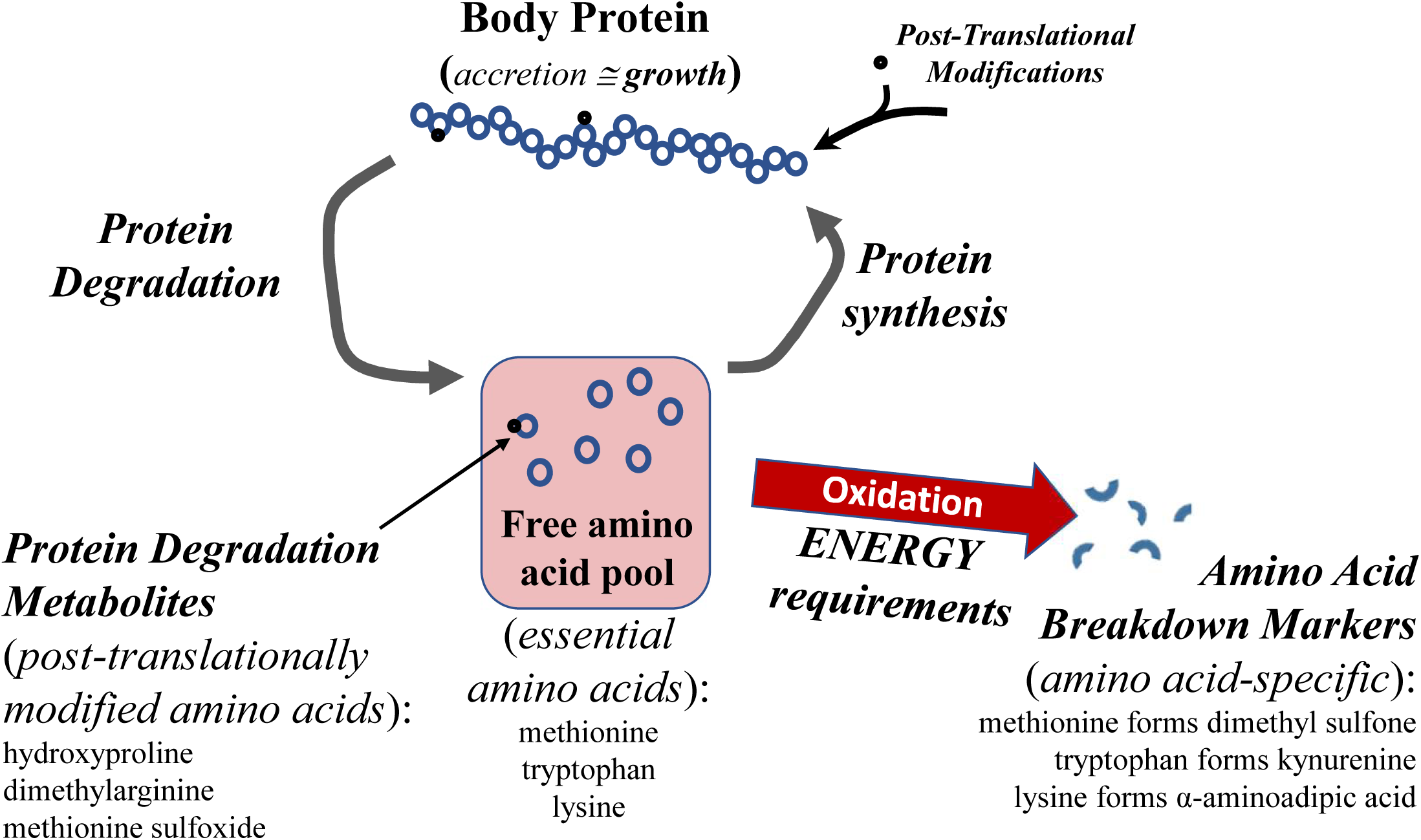
Conceptual diagram linking protein degradation and markers of amino acid breakdown to walleye (*Sander vitreus*) growth, diet, and energy requirements. Throughout this manuscript, amino acid breakdown refers to breaking down an amino acid to use it as an energy source. Metabolites specific to the breakdown pathways of specific essential amino acids, indicated below, were selected as markers of these processes. Meanwhile protein degradation refers to proteins that are being separated back into their constituent amino acids. For the current study, we used metabolites in the blood that would come specifically from post-translationally modified amino acids because their release is the result or proteins that have been synthesized and subsequently degraded.

Generally, if fish are not taking in sufficient food, stored lipids and carbohydrates are preferentially used to meet energetic demands, with whole-body protein being increasingly mobilized as a fuel after lipid and carbohydrate reserves are depleted (Black and Love 1986; Collins and Anderson 1995). Essential, or non-dispensable, amino acids may be useful for insight into amino acid oxidation in wild fish because the animal must get them through its diet, and cannot rely solely on its own metabolic processes for their supply. If dietary essential amino acid intake is insufficient, the fish cannot grow or may even fail to maintain body mass. For this reason, increased breakdown of essential amino acids would coincide with either excess intake from the diet or a need to catabolize body protein to make up for an energetic deficiency. Amino acid breakdown can be implicated by the presence of metabolites that result from their breakdown, and essential amino acids (specifically methionine, tryptophan, and lysine in the current study) are of particular use in this context because their presence represents only food intake or protein breakdown. Assuming conserved pathways of amino acid breakdown across animals, methionine breakdown results in the formation of dimethyl sulfone (Engelke et al., 2005), tryptophan is linked to kynurenine production (Knox and Mehler, 1950), and lysine to α-aminoadipic acid (Borsook et al., 1948). As such, we argue that the elevation essential amino acids and concomitant increase in levels of their amino acid-specific breakdown products may indicate a physiological need for fish to increase the oxidation of these specific amino acids for energy, over preferential recycling of essential amino acids back into protein.

To assess possible biomarkers of protein degradation, our strategy focused on ‘post-translational’ protein modifications, which are physical changes to amino acids following their incorporation into a protein molecule. These post-translational modifications are in some cases retained by the amino acid following degradation of the protein back to free amino acids (Figure 2). Thus the presence of these modified amino acids in blood may act as potential markers of protein breakdown. Here, we focus on three modified amino acids: hydroxyproline, a major constituent of collagen (Stetten, 1949; Prockop and Sjoerdsma, 1961), dimethylarginines, which are enzymatic modifications of arginine (Vallance and Leiper, 2004), and methionine sulfoxide, which is a non-enzymatic modification of methionine that can occur in 4-8% of all methionine found within proteins (Stadtman et al., 2005). The proteinaceous origin of these post-translationally modified metabolites may thus elucidate the relative rate of protein degradation in walleye from different areas of Lake Winnipeg (see Figure 2).

### Linking metabolites and spatial patterns

In the present study, we examined both temporal and spatial changes in Lake Winnipeg walleye morphological measurements using gillnet index data collected by the Province of Manitoba. First, length-at-age was used to assess longer-term trends in growth prior to 2017. Differences in walleye length-at-age may be an effective metric to study the historical rainbow smelt collapse (approximately 2013), since length-at-age is a summary of prior growth while nutritional deprivation has little effect on length (Caruso et al., 2011; Einen and Thomassen, 1998; Sumpter et al., 1991). On the other hand, body condition or length-mass relationships can vary with a single spawning or feeding event, each of which change a fish’s mass. Spatial patterns in length-at-age (≈ growth rate) and length-mass relationships (≈ body condition) are thus assessed concurrently. We then linked length-mass relationships to a survey of walleye in Lake Winnipeg in 2017, where blood was sampled non-lethally for metabolites included in our nutritional biomarker development strategy (see above; Figure 2). Spatial differences in the presence of chosen metabolites were then tested with estimated marginal means (Lenth, 2019; Searle et al., 1980) and linear models incorporating length and site collected. We hypothesized that because the rainbow smelt collapse was most pronounced in the north basin of Lake Winnipeg (Manitoba Government, 2018), north basin walleye would exhibit the greatest decline in relative mass over time and length-at-age. We also hypothesized that metabolites would implicate higher rates of endogenous protein breakdown (i.e., elevated hydroxyproline, dimethylarginine, and methionine sulfoxide) and amino acid oxidation (i.e., elevated dimethyl sulfone, kynurenine, and α-aminoadipic acid) in the north basin than in the south basin. This work thus 1) explored the effect of the rainbow smelt collapse on Lake Winnipeg walleye morphology and 2) examined the potential for a suite of 9 metabolites to reflect walleye nutritional status and be developed into a non-lethal sampling approach for assessing the nutritional state of wild-caught fish.

## Methods

### Lake Winnipeg Gillnet Index Data

We used the Lake Winnipeg gillnet index data collected by the Government of Manitoba between 2009 and 2018 (data accessed September 2019: https://www.gov.mb.ca/sd/fish_and_wildlife/fish/commercial_fishing/netting_data.html). This gillnet index has been run annually by the province since 1979 to provide an alternative to commercial fisheries data to better track trends in size and abundance for walleye and sauger (*Sander canadensis*) in Lake Winnipeg. We focused on the data collected between 2009 and 2018 because there was consistent monitoring of the same six sites during that period and these data included age estimates for each fish. To examine walleye length-mass and length-at-age relationships over time, sauger and dwarf walleye (which possibly inhabit a different ecological niche and are responding to selective pressures; see Moles et al., 2010) were filtered out. The remaining data set included year, site, basin collected, gillnet mesh size, fork length, mass, and sex for adult and sub-adult walleye. Fork length and mass were transformed on a log_10_ scale. Gillnet mesh size was included as an independent variable where possible to account for bias in walleye length and mass caught in gillnets with different mesh sizes. For all linear models, we chose variables based on their biological significance, as opposed to taking a stepwise model selection approach. Statistical modeling was performed in R (R Core Team, 2019) using the packages emmeans (Searle et al., 1980), sjstats (Lüdecke, 2019), and tidyverse (Wickham et al., 2019). Scripts used for all statistical analyses in this study are available at: https://github.com/BioMatt/walleye_condition_metabolites.

### Lake Winnipeg Walleye Length-at-Age Over Time

Changes in growth rate over time were assessed for walleye from seven sites across Lake Winnipeg using length-at-age data from the gillnet index, with a special focus on growth rates before and after the rainbow smelt collapse (in approximately 2013). Length is a useful measure because a fish facing nutritional deprivation decreases in mass, but decreases in length either negligibly or not at all (Caruso et al., 2011; Einen and Thomassen, 1998; Sumpter et al., 1991). Length-at-age therefore represents more long-term trends in growth than year-specific values for body condition or length-mass relationships. A separate linear model was used for each site sampled for the gillnet index data, with Dauphin River and Grand Rapids representing the north basin, Matheson Island and Frog Bay representing the channel, and Riverton and Grand Beach representing the south basin of Lake Winnipeg (Figure 1). These linear models used fork length as the dependent variable, and the independent variables were age interacting with year, sex, and gillnet mesh size. Results were therefore averaged over gillnet mesh sizes to account for sampling bias. Only walleye ages two through six years old as estimated by counting annuli in otoliths were included in the models, because not all sites had sufficient sample sizes available for walleye of older and younger ages (*n* ≥ 10 individuals age class^−1^ year^−1^ site^−1^, except the Dauphin River age two fish in 2012 and 2014 were included, *n*=5 and 2, respectively). A total of *n*=2164 individuals were used in the models for Dauphin River, *n*=2745 for Grand Rapids, *n*=844 for Matheson Island, *n*=959 for Frog Bay, *n*=1732 for Riverton, and *n*=1729 for Grand Beach collection sites.

Differences in walleye growth rates (i.e., length-at-age) among sampling sites were also examined for the years surrounding 2017 (i.e., 2015–2018), to support connections with the available metabolite data (i.e., from year 2017) and thus potential food availability during this time. Age two and three walleye sampled in 2018 were of particular interest for this analysis because length-at-age for young walleye in the growth phase of their life cycle in 2018 should reflect food availability from 2017 through to their capture in 2018. Four similar linear models (i.e., one for each year) were used for length-at-age comparisons among sites. In each model, fork length was the dependent variable, while independent variables were the interaction of site and age, sex, and gillnet mesh size. As above, only walleye ages two through six years old were included in the models. For these analyses, *n*=1844 individuals were available for the year 2015, *n*=1392 for 2016, *n*=805 for 2017, and *n*=624 for 2018. Estimated marginal means and estimated marginal trends were used to investigate pairwise mean and slope differences between sites for each model (Lenth, 2019; Searle et al., 1979). Briefly, the analysis of estimated marginal means obtains predictions from a linear model using Tukey’s post hoc test and finds meaningful averages to summarize primary factor effects while estimated marginal trends follows the same process but for the interaction between two predictors in a model (see Lenth, 2019).

### Lake Winnipeg Walleye Relative Mass Over Time

Relative mass over time was examined for walleye across Lake Winnipeg. These relative mass measures were used as metrics analogous to body condition, which is an imperfect approach to our data because walleye mass may change in a single feeding or spawning event, thus changing a fish’s body condition and length-mass relationship. However, an examination of relative mass remains useful because it represents a link between inter-annual growth rate trends and metabolite presence, which can vary on much faster timescales. Only walleye ≥375 mm in fork length from the gillnet index data, *n*=5,838 individuals (mean 583.80 walleye ± 220.17 s.d. year^−1^), were included in the model because this value represents the smallest estimate of fork length for mature individuals among Manitoba lakes (Craig et al., 1995), and is a conservative threshold for modeling the fish that were sampled for metabolites (minimum fork length 452 mm). After filtering, a total of *n*=5,838 individuals (mean 583.80 walleye ± 220.17 s.d. year^−1^) remained.

To study how relative mass has changed over time, a linear model was used with mass as the dependent variable, and independent variables were fork length and its interactions with site, year and sex. Eta squared is reported for the independent variables in this model, where analysis of variance applied to the model outputs returns the percentage of variance in the final model accounted for by an independent variable (Levine and Hullett, 2002).

### Spatial Differences Among Walleye in 2017

In addition to exploring length-mass relationships over time, we modeled spatial relationships among Lake Winnipeg walleye using the gillnet index data set in 2017 to establish possible connections between length-mass relationships and metabolite presence in that year. Here, *n*=286, 127, and 227 individuals remained from the Dauphin River, Matheson Island, and Riverton sites, respectively. These three sites were chosen because they are most similar to the three sites sampled for metabolites (i.e. Dauphin River, Matheson Island, and the Red River).

This model used mass as the dependent variable, with the independent variables as the interaction of mass with fork length, site collected, sex, age, and gillnet mesh size.

Another linear model was used to assess the length-mass relationship of walleye caught as part of the blood metabolite study in 2017. The smallest walleye in this data set was 452 mm in fork length. Mass was the dependent variable, with the independent variables fork length interacting with site collected, sex, and the interaction of fork length and sex. Note that gillnet mesh size and age were not included in the model since fish were caught by electrofishing and age was not known for these fish.

To examine how similar the length-mass relationships were of walleye were between those collected for metabolite analysis and those for the gill net index data, we used a linear model with an independent variable describing which study an individual was collected for. Here, mass was the dependent variable, which was related to fork length, basin collected, sex, study (metabolite or gillnet index), the interaction of fork length and basin, the interaction of fork length and sex, and the interaction between fork length and study. In this model, only gillnet index samples from the Dauphin River, Matheson Island, and Riverton were included. Riverton from the gillnet index data and the Red River from the metabolite data were used to jointly represent the south basin of Lake Winnipeg.

### Metabolite Data Collection

In 2017, 39 walleye were collected from three sampling locations: the Red River, Matheson Island and Dauphin River representing the south basin, channel, and north basin of Lake Winnipeg, respectively (Figure 1). Each sampling location was at a known spawning site during spawning season (May 2^nd^ in the Red River, May 17^th^ and 18^th^ at Matheson Island, and May 29^th^ through 31^st^ at the Dauphin River). Measured metabolites may thus have been affected by the approximately one-month range in sampling. In addition, we could not verify that sampled individuals had spawned at their collection site, or if they had spawned elsewhere and moved to the collection site. Both males and females were collected, with *n*=17 from the Red River (2 males, 15 females), *n*=5 from Matheson island (2 males, 3 females), and *n*=17 from the Dauphin River (5 males, 12 females), and had a minimum mass of 1 kilogram (mean 2.32 kg ± 0.97 s.d.). Individuals were collected by boat electrofishing, held in a live well for no longer than one hour, and anaesthetized using a Portable Electroanesthesia System (PES™, Smith Root, Vancouver, Washington, USA) in accordance with approved animal use protocols of Fisheries and Oceans Canada (FWI-ACC-2017-001, FWI-ACC-2018-001), the University of Manitoba (F2018-019) and the University of Nebraska-Lincoln (Project ID: 1208).

One milliliter of whole blood was collected for metabolite analysis from anaesthetized walleye by caudal puncture using a heparinized needle and 3 ml syringe, flash frozen in liquid nitrogen immediately after sampling, and stored at −80°C. Of note, logistics prevented separation of plasma or serum and therefore all metabolite data are for whole blood. This effort was part of a larger study assessing the physiological health, movement, and genetic structure of walleye in Lake Winnipeg. Other tissues collected at the time of blood sampling include a fin clip, gill filaments, the first dorsal spine, scales, and a muscle biopsy. All fish were sampled non-lethally and a VEMCO acoustic tag (VEMCO, Bedford, Nova Scotia, Canada) was surgically implanted prior to release back into the water near the collection site.

### Blood Sample Analyses

Walleye blood samples collected in 2017 (see above) were analyzed using nuclear magnetic resonance (NMR) spectroscopy and a combination of direct injection mass spectrometry with a reverse-phase liquid chromatography with mass spectrometry (DI/LC-MS/MS) assay at the University of Alberta Metabolomics Centre TMIC (Edmonton, AB, Canada) as part of a large scale targeted metabolic study. Here we examine a small subset of analytes to identify potential biomarkers of protein degradation and amino acid breakdown as described in Figure 2. Both NMR and DI/LC-MS/MS detected methionine and lysine, so measurements for each metabolite were averaged over the two detection methods. Tryptophan and dimethyl sulfone were only measured by NMR. Kynurenine, hydroxyproline, dimethyl sulfone, and α-aminoadipic acid were detected with DI/LC-MS/MS.

For NMR spectroscopy, deproteinization, involving ultra-filtration, was performed to remove proteins. Filtration, and centrifugation steps were subsequently done to further purify the sample (Psychogios et al., 2011). 250 µL of the blood sample was transferred to a 3 mm SampleJet NMR tube for subsequent spectral analysis following a protocol based on Saude et al. (2004). NMR spectra was collected on a 700 MHz Avance III (Bruker) spectrometer and the spectra acquired at 25°C. NMR spectra were processed and analyzed using the Chenomx NMR Suite Professional software package version 8.1 (Chenomx Inc., Edmonton, AB). DI**/**LC-MS/MS was done on an API4000 Qtrap® tandem mass spectrometry instrument (Applied Biosystems/MDS Analytical Technologies, Foster City, CA) equipped with an Agilent 1260 series HPLC system (Agilent Technologies, Palo Alto, CA). The samples were delivered to the mass spectrometer by a LC method followed by a direct injection (DI) method. Data analysis was done using Analyst 1.6.2.

### Modeling Metabolites Differences

Differences in nine metabolites across sampling sites were assessed using separate linear models. Dimethyl sulfone, kynurenine, α-aminoadipic acid, methionine, tryptophan, lysine, hydroxyproline, dimethylarginine, and methionine sulfoxide were used as dependent variables in their respective linear models. For each model, the independent variables were log fork length and site collected. Estimated marginal means of metabolite presence were calculated to establish pairwise significance between sites. Significance for each fork length and site collected were calculated, along with eta squared (Levine and Hullett, 2002) to report effect size for each independent variable.

## Results

### Lake Winnipeg Walleye Length-at-Age Over Time

Each of the Dauphin River, Grand Rapids, Matheson Island, Frog Bay, Riverton, and Grand Beach sites exhibited a decline in length-at-age for age six walleye between the years 2012 and 2018 (Figure 3). However, for the age two and three walleye, while the Dauphin River (north basin) site also showed a shorter length-at-age in 2018 compared to 2012, the south basin sites (Riverton and Grand Beach) showed similar length-at-age in later years (i.e. 2017 and 2018) relative to earlier years (i.e. 2012 and 2013). The Grand Rapids (in the north basin) showed the most consistent decline in length-at-age for all ages over time except for the age two fish sampled in 2017 and 2018 (Figure 3).

**Figure 3.**
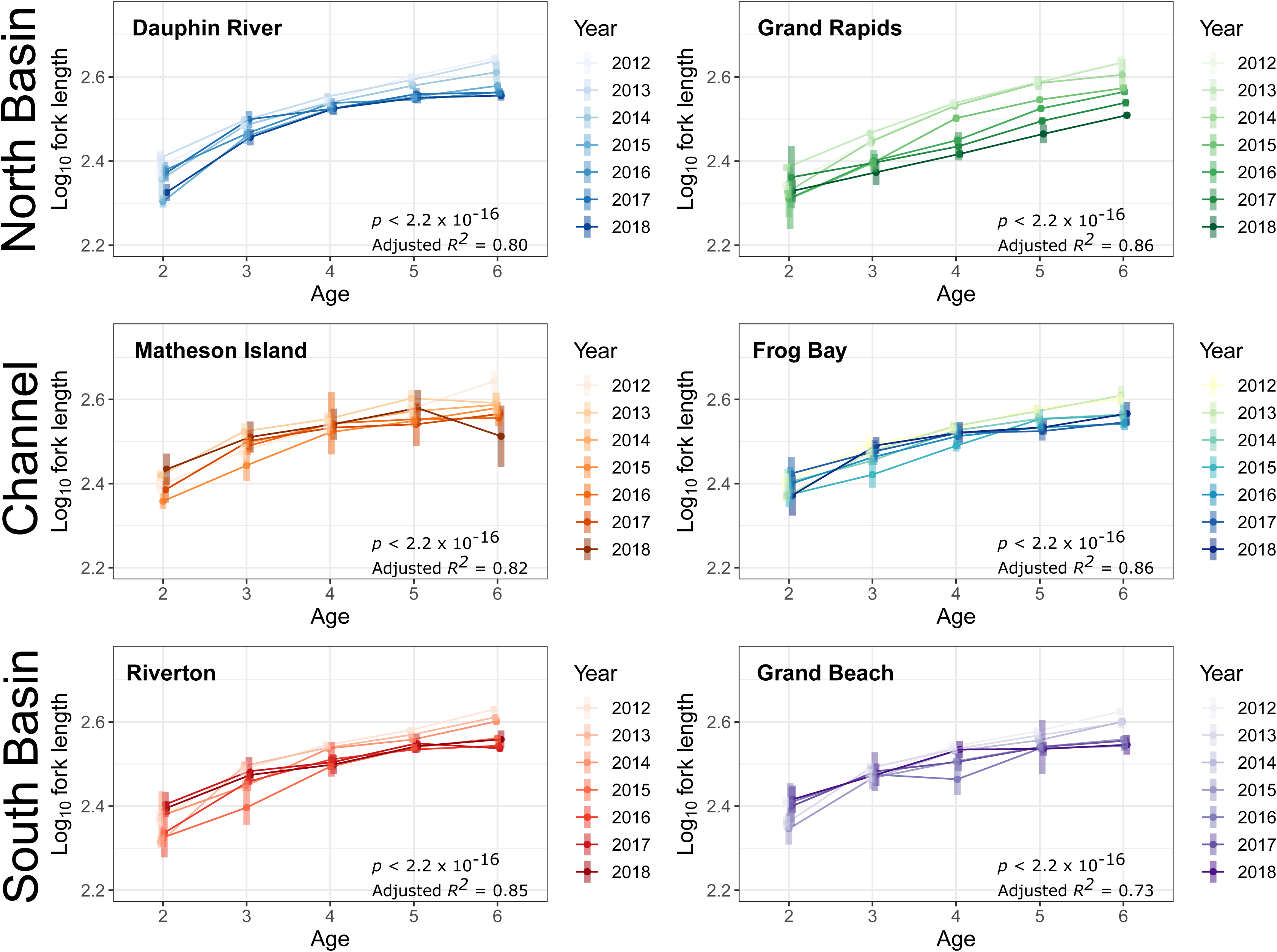
Estimated marginal means of log_10_ fork length-at-age for walleye (*Sander vitreus*) from 2012 to 2018 across gillnet index collection sites in Lake Winnipeg. The data were collected by the Government of Manitoba. These figures are derived from linear models relating log_10_ fork length to the interaction of age with year, sex, and gillnet mesh size. A separate model was used for each collection site. 95% confidence intervals are provided as error bars at each age. Overall model significance and adjusted *R*^2^ is provided in each panel.

Length-at-age decreased for Dauphin River walleye at ages two and three between the years 2017 and 2018 (Figure 3, Figure 4). For length-at-age in 2018 specifically, pairwise estimated marginal means using Tukey’s post-hoc test on the linear model (*F* = 66.81, *p* < 2.2 × 10^−16^, adjusted *R^2^* = 0.80) showed significant differences in estimated marginal mean of length-at-age between the Dauphin River (north basin) and Riverton (south basin), Grand Beach (south basin), and Matheson Island (channel) sites in age two walleye (Table 1). Slopes in length-at-age relationships, as described using estimated marginal trends, were also significantly more steep between the Dauphin River and the Matheson Island, Riverton, and Grand Beach sites, respectively (Table 1, Figure 4). Based on these results, we suggest that young Dauphin River walleye grew more slowly in 2017 than young walleye at the Matheson Island (channel) and the south basin sites.

**Table 1.** *P*-values of pairwise estimated marginal means and trends calculated using Tukey’s post hoc test, pulled from a linear model relating log_10_ fork length to age in interaction with site, sex, and mesh size in the year 2018 (*F* = 66.81, *p* < 2.2 × 10^−16^, adjusted *R*^2^ = 0.80). Means, representing differences in intercepts are above the diagonal and trends, representing differences in slopes are below the diagonal. Estimated marginal means are specific to age two walleye (*Sander vitreus*), while estimated marginal trends are calculated across all ages between two and six. Data included in this model are from the gillnet index collected by the Government of Manitoba.

|  | Grand Rapids | Dauphin River | Matheson Island | Frog Bay | Riverton | Grand Beach |
| --- | --- | --- | --- | --- | --- | --- |
| Grand Rapids | - | 0.67 | < 0.05 | 0.24 | < 0.05 | < 0.05 |
| Dauphin River | 0.0015 | - | 0.0003 | 0.70 | < 0.05 | 0.0003 |
| Matheson Island | 0.96 | 0.0035 | - | 0.61 | 0.85 | 1.0 |
| Frog Bay | 0.69 | 0.96 | 0.40 | - | 0.94 | 0.61 |
| Riverton | 1.0 | 0.0002 | 0.94 | 0.67 | - | 0.85 |
| Grand Beach | 0.92 | 0.0005 | 1.0 | 0.30 | 0.86 | - |

**Figure 4.**
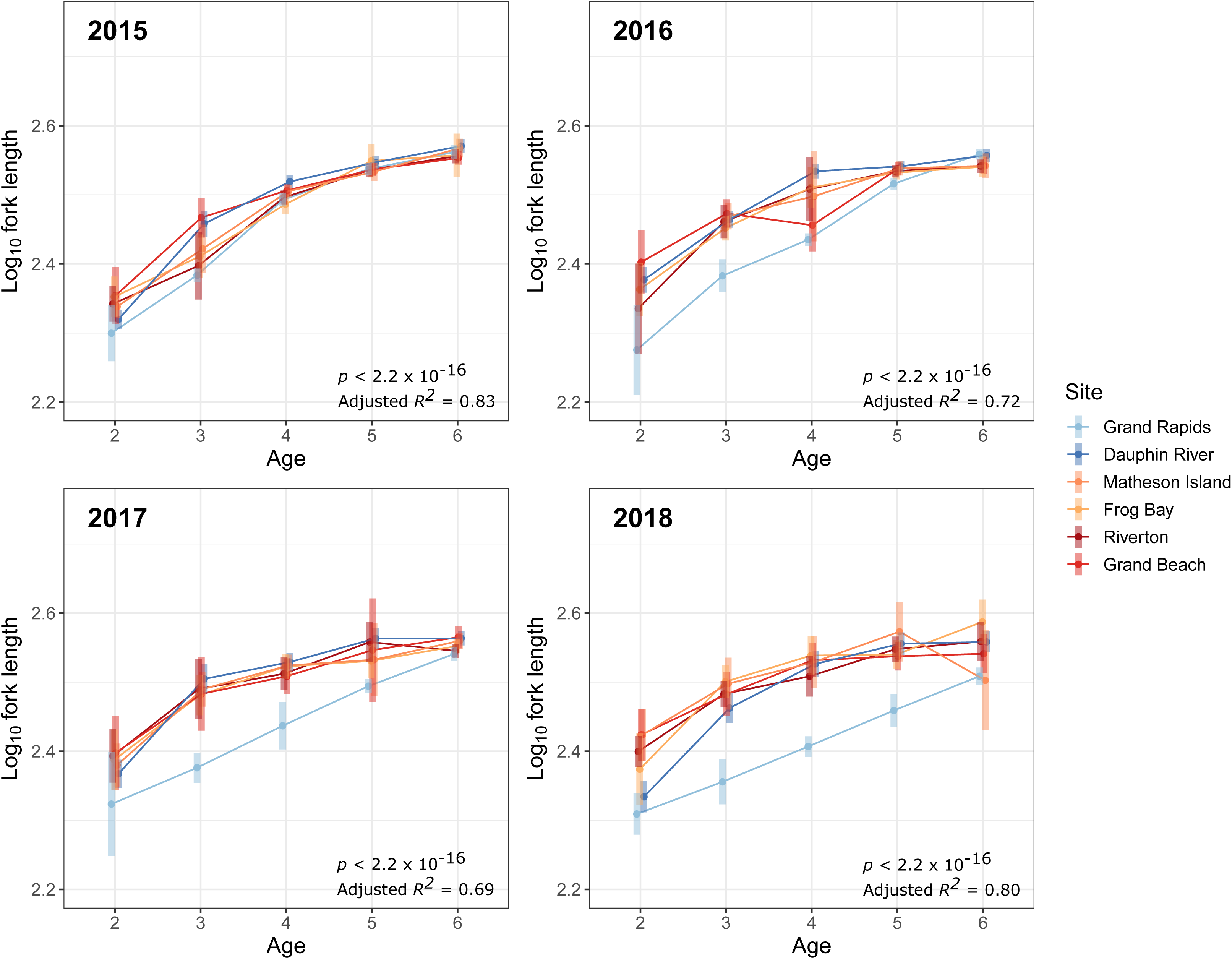
Estimated marginal means of log fork length-at-age between 2015 and 2018 across all sites collected for the gillnet index data in Lake Winnipeg walleye (*Sander vitreus*). The data were collected by the Government of Manitoba. These figures are derived from linear models relating log fork length to age in interaction with site, sex, and mesh size. A separate linear model was used for each of the years 2015, 2016, 2017, and 2018. 95% confidence intervals are provided as error bars at each age. Overall model significance and adjusted *R*^2^ is provided in each panel.

### Lake Winnipeg Walleye Relative Mass Over Time

From the years 2009 through 2018, year, sex, and the interaction between fork length and year had a significant relationship with Lake Winnipeg walleye length-mass relationships (*F* = 2899, *p* < 2.2 × 10^−16^, adjusted *R*^2^ = 0.95, Table 2). While fork length had the greatest effect size as indicated by eta squared, year had a greater effect size than site collected during the time period studied. Confidence intervals determined using estimated marginal means indicated significant differences among the effect of years on mass, with a drop in predicted mass most noticeable between the years 2014 and 2015 (Figure 5A).

**Table 2.** Results from a linear model describing relative mass over time, relating log_10_ mass to log_10_ fork length and its interaction with year, site, and sex, with mesh size controlled for (*F* = 2899, *p* < 2.2 × 10^−16^, *R*^2^ = 0.95). The gillnet index data between the years 2009 and 2018 collected by the Government of Manitoba were used for this length-mass model. Only walleye (*Sander vitreus*) ≥375 millimeters in fork length are included.

|  | *p*-value | eta squared |
| --- | --- | --- |
| Log<sub>10</sub> Fork Length | < 0.05 | 0.25 |
| Year | < 0.05 | 0.0080 |
| Site | 0.11 | 0.0010 |
| Sex | 0.040 | 0.0 |
| Mesh Size | < 0.05 | 0.020 |
| Log<sub>10</sub> Fork Length * Year | < 0.05 | 0.0070 |
| Log<sub>10</sub> Fork Length * Site | 0.16 | 0.0010 |
| Log<sub>10</sub> Fork Length * Sex | 0.053 | 0.0 |

**Figure 5.**
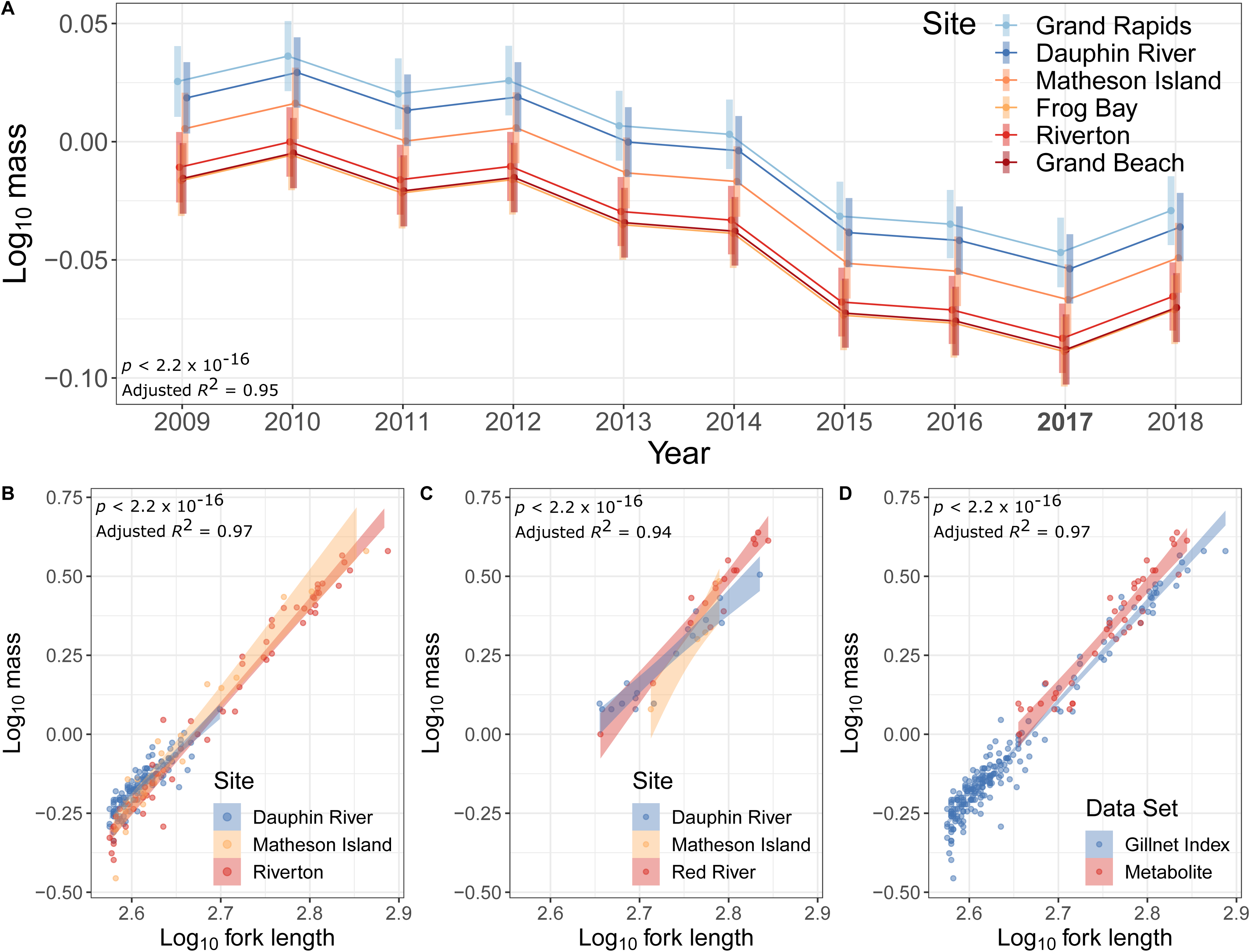
Relative mass over time and length-mass relationships among Lake Winnipeg walleye (*Sander vitreus*) caught in 2017. Panel A represents estimated marginal means of log_10_ mass between 2009 and 2018 across all sites collected for the gill net index data in Lake Winnipeg walleye (*Sander vitreus*). These data are pulled from a linear model relating log_10_ mass as a dependent variable to log_10_ fork length, sex, site, and mesh size as independent variables (*F* = 2686, *p* < 2.2 × 10^−16^, adjusted *R*^2^ = 0.98). 95% confidence intervals are provided as error bars at each year. For panel B, the linear model related log_10_ mass to the interaction of log_10_ fork length and basin collected, sex, mesh size, and age (*F* = 222, *p* < 2.2 × 10^−16^, adjusted *R*^2^ = 0.97) using the gillnet index data from the Dauphin River, Matheson Island, and Riverton sites. Panel C represents a linear model on walleye used for fish from the metabolite measurements, and this linear model related log_10_ mass in kg to log_10_ fork length interacting with site collected, sex, and the interaction of log_10_ fork length and sex (*F* = 86, *p* < 2.2 × 10^−16^, adjusted *R*^2^ = 0.94). Panel D shows a linear model comparing length-mass relationships in walleye between those used for metabolites and from the gillnet index data. This model relates log_10_ mass as the dependent variable, with log_10_ fork length, basin collected, sex, study (metabolite or gill net index), the interaction of log_10_ fork length and basin, the interaction of log_10_ fork length and sex, and the interaction between log_10_ fork length and study as independent variables (*F* = 889, *p* < 2.2 × 10^−16^, adjusted *R^2^* = 0.97). Standard error is shown with filled colors. Overall model significance and adjusted *R*^2^ is provided in each panel. The gillnet index data were collected by the Government of Manitoba and only walleye ≥375 millimeters in fork length are included in this models, while in the metabolite data are large fish (≥452 mm in fork length) caught by electrofishing, from which whole blood metabolites were measured. Log_10_ mass was calculated in kilograms, and log_10_ fork length in millimeters.

### Spatial Differences Among Walleye in 2017

In a linear model using gillnet index data for fish captured in 2017 that were ≥375 mm in fork length and including only the Dauphin River, Matheson Island, and Riverton sites, (*F* = 222, *p* < 2.2 × 10^−16^, adjusted *R*^2^ = 0.97), estimated marginal mean length-mass relationships were the same for three sites, but estimated marginal trends were lower in the Dauphin River compared to Matheson Island (Table 3; Figure 5B). This suggests a different length-mass relationship for the north basin fish compared to walleye caught in the channel.

**Table 3.** *P*-values of pairwise estimated marginal means and trends between either basins or sites for two different linear models. One linear model uses walleye (*Sander vitreus*) ≥375 mm in fork length from the gillnet index and relates log_10_ mass to log_10_ fork length and site collected, sex, gillnet mesh size, and age (*F* = 222, *p* < 2.2 × 10^−16^, adjusted *R*^2^ = 0.97). Data included in this model are from the gillnet index collected by the Government of Manitoba. The other linear model uses data from walleye sampled for metabolites (≥452 mm in fork length), and relates log_10_ mass to the interaction of log_10_ fork length and site collected, sex, and the interaction of log_10_ fork length and sex (*F* = 86, *p* < 2.2 × 10^−16^, adjusted *R*^2^ = 0.94). *P*-values for means, representing differences in intercepts are above the diagonal and *p*-values for trends, representing differences in slopes are below the diagonal.

|  | Gill Net Index |  |  | Metabolite |  |  |
| --- | --- | --- | --- | --- | --- | --- |
|  | Dauphin River | Matheson Island | Riverton | Dauphin River | Matheson Island | Red River |
| Dauphin River | - | 0.60 | 0.082 | - | 0.65 | 0.081 |
| Matheson Island | 0.024 | - | 0.60 | 0.037 | - | 0.040 |
| Riverton/Red River | 0.16 | 0.41 | - | 0.060 | 0.35 | - |

Similar to the larger fish sampled as part of the gill net index in 2017, for length-mass relationships among walleye collected for metabolites (*F* = 86, *p* < 2.2 × 10^−16^, adjusted *R*^2^ = 0.94), estimated marginal trends were also lower in the Dauphin River compared to Matheson Island (Table 3; Figure 5C). However, unlike the linear model used with the gillnet index data, the Red River (south basin) showed a higher estimated marginal mean mass than the Matheson Island (channel) site (Table 3; Figure 5C).

When comparing length-mass relationship of larger walleye collected in 2017 as part of the gill net index and metabolite studies, no significant effect of study and of study interacting fork length was found (*F* = 889, *p* < 2.2 × 10^−16^, adjusted *R*^2^ = 0.97, Table 4). However, estimated marginal means of mass based on fork length were higher in the walleye from the metabolite data (*p* = 0.00070) while estimated marginal trends for fork length-mass relationships between studies were not different (*p* = 0.90). A plot of fork length and mass with these data reveals that the metabolite-measured walleye lie at the upper end of the distribution in length-mass for walleye collected for the gillnet index data, while following a similar slope (Figure 5D).

**Table 4.** Results from a linear model relating log_10_ mass to the log_10_ fork length, basin collected, study of data origin (gillnet index or metabolite), sex, the interaction of log_10_ fork length and basin, and the interaction of log_10_ fork length and study (*F* = 889, *p* < 2.2 × 10^−16^, *R*^2^ = 0.97). This model represents the length-mass relationship for walleye (*Sander vitreus*) collected for the gillnet index data by the Government of Manitoba and for metabolite information by the authors. These walleye were collected in 2017, and only fish ≥375 millimeters in fork length are included in this model. From the gillnet index data, only walleye from the Dauphin River, Matheson Island, and Riverton sites are included.

| | $p$-value | eta squared |
| --- | --- | --- |
| Log<sub>10</sub> Fork Length | < 0.05 | 0.74 |
| Basin | 0.0010 | 0.020 |
| Study | 0.98 | 0.0 |
| Sex | 0.65 | 0.0 |
| Log<sub>10</sub> Fork Length * Sex | 0.62 | 0.0 |
| Log<sub>10</sub> Fork Length * Basin | < 0.05 | 0.019 |
| Log<sub>10</sub> Fork Length * Study | 0.90 | 0.0 |

### Modeling Metabolites Differences

Metabolite presence of the three essential amino acids varied significantly across sampling sites for Lake Winnipeg walleye sampled in 2017. The models predicting the presence of each of the three free essential amino acids were each significant (*F* = 4.3, 4.7, and 3.7, *p* = 0.011, 0.0071, and 0.020, adjusted *R*^2^ = 0.21, 0.23, and 0.18 for methionine, tryptophan, and lysine respectively, Table 5). In addition, site was a significant independent variable within the overall models for the three essential amino acids while fork length was not, with eta squared higher for site than for fork length in each case (Table 5). Estimated marginal means for each of the three essential amino acids revealed significant differences in predicted essential amino acid presence based on site, with higher values in the Dauphin River (north basin) than in the Red River (south basin) (Figure 6A, B, and C).

**Table 5.** Results from linear models relating metabolite presence to log_10_ fork length and site collected, with a separate model for each metabolite. Category represents the conceptual framework used to classify the nine metabolites studied (see Figure 2 for details). Overall model *p*-values are provided under their respective metabolites. *P*-values and eta squared are reported for log_10_ fork length and site collected as independent variables within models. Briefly, methionine, tryptophan, and lysine represent essential amino acids. Dimethyl sulfone, kynurenine, and α-aminoadipic acid represent essential amino acid breakdown. Last, hydroxyproline, dimethylarginine, and methionine sulfoxide represent endogenous protein breakdown. The walleye (*Sander vitreus*) from which metabolites were measured for these linear models were *n*=39 individuals (≥452 mm in fork length) collected by boat electrofishing in 2017 from the Dauphin River, Matheson Island, and Red River representing the north basin, channel, and south basin of Lake Winnipeg, respectively. Metabolites were measured from whole blood.

| Category | Amino acid | Independent variable | *p*-value | eta squared |
| --- | --- | --- | --- | --- |
| Essential Amino Acids | Methionine | Log<sub>10</sub> fork length | 0.74 | 0.0020 |
|  | *p* = 0.011 | Site collected | 0.0070 | 0.25 |
|  | Tryptophan | Log<sub>10</sub> fork length | 0.36 | 0.019 |
|  | *p* = 0.0071 | Site collected | 0.032 | 0.17 |
|  | Lysine | Log<sub>10</sub> fork length | 0.52 | 0.010 |
|  | *p* = 0.020 | Site collected | 0.046 | 0.16 |
| Amino Acid Breakdown Markers | Dimethyl Sulfone | Log<sub>10</sub> fork length | 0.088 | 0.070 |
|  | *p* = 0.031 | Site collected | 0.36 | 0.048 |
|  | Kynurenine | Log<sub>10</sub> fork length | 0.80 | 0.002 |
|  | *p* = 0.070 | Site collected | 0.092 | 0.13 |
| | α-Aminoadipic Acid | Log<sub>10</sub> fork length | 0.94 | 0.0 |
|  | *p* = 0.81 | Site collected | 0.66 | 0.023 |
| Protein Degradation Metabolites | Hydroxyproline | Log<sub>10</sub> fork length | 0.69 | 0.002 |
| | *p* = $1.3 \times 10^{-18}$ | Site collected | $< 2.2 \times 10^{-16}$ | 0.60 |
|  | Dimethylarginine | Log<sub>10</sub> fork length | 0.97 | 0.0 |
|  | *p* = 0.0017 | Site collected | 0.0020 | 0.30 |
|  | Methionine Sulfoxide | Log<sub>10</sub> fork length | 0.47 | 0.011 |
|  | *p* = 0.0016 | Site collected | 0.0080 | 0.24 |

**Figure 6.**
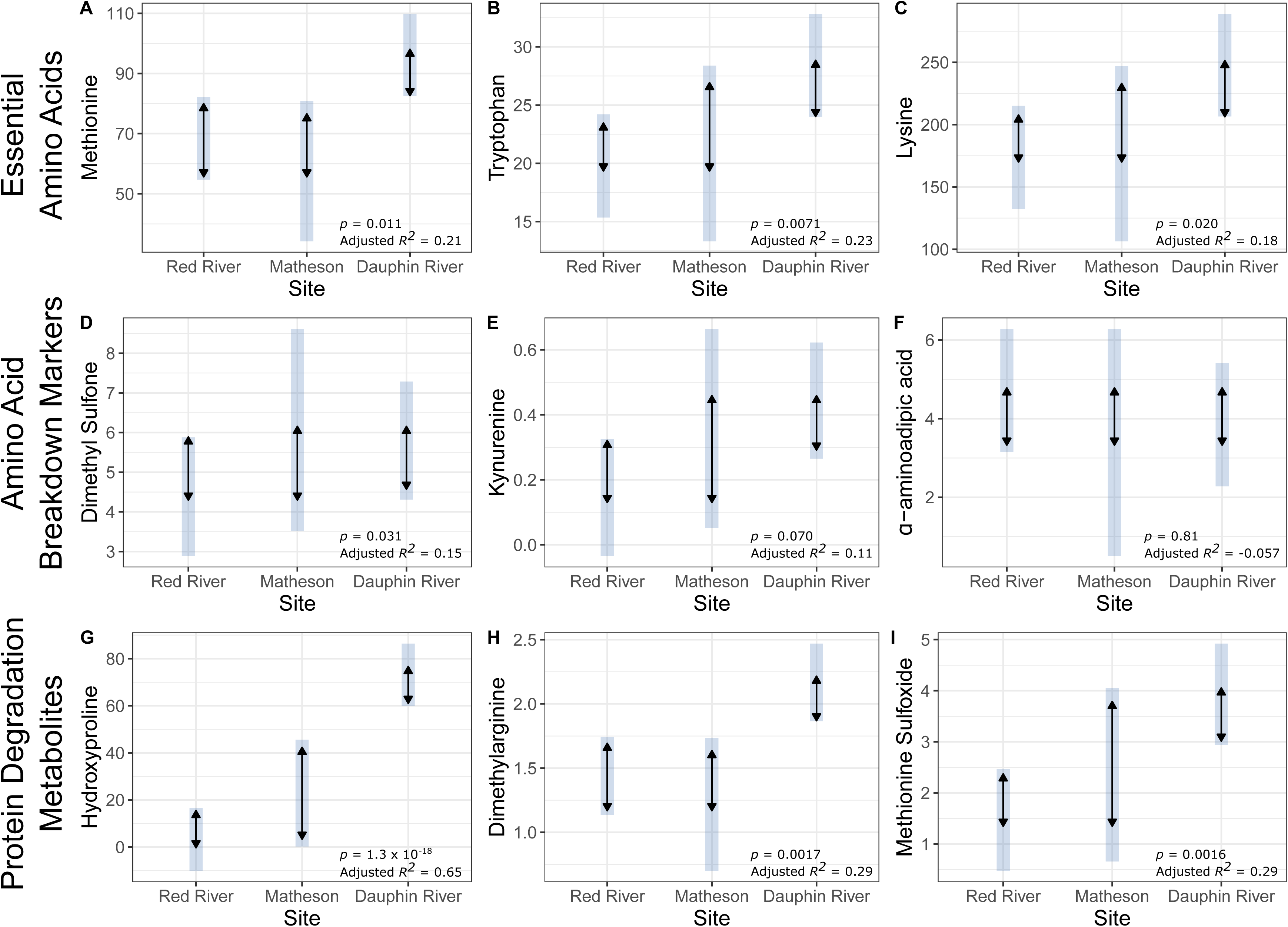
Estimated metabolite concentration (µmol l^−1^ in whole blood) for Lake Winnipeg walleye (*Sander vitreus*) by site, predicted by linear models that incorporate metabolite as the dependent variable, with log_10_ fork length and site collected as the independent variables. The three sites on the horizontal axis are the Red River (south basin), Matheson Island (channel), and Dauphin River (north basin). Here, *n*=39 large fish (≥452 mm in fork length) were caught by electrofishing in 2017. On the vertical axis are estimated marginal means for each metabolite. The light blue bars represent 95% confidence intervals, while the black arrows represent significance for pairwise comparisons between sites. Overall linear model significance and adjusted *R*^2^ is provided in each panel. Panels A, B, and C represent essential amino acids. Panels D, E, and F represent respective amino acid breakdown metabolites that in conjunction with plots A, B, and C, may describe amino acid oxidation. Panels G, H, and I represent metabolites associated post-translational modification and therefore may reflect endogenous protein degradation (see Figure 2 for additional details).

Overall, there was no effect of sampling location or fish size (i.e., fork length) on essential amino acid breakdown biomarkers. Linear models for α-aminoadipic acid, kynurenine, and dimethyl sulfone were not significant (*F* = 0.32, 2.6, and 3.3, *p* = 0.81, 0.070, and 0.031, adjusted *R*^2^ = 0.057, 0.11, and 0.15, respectively, Table 5), indicating a failure to fit fork length and site to essential amino acid breakdown metabolite presence. The model for dimethyl sulfone was significant (*p* = 0.031), but neither site nor fork length were significant independent variables (*p* = 0.36 and 0.088, respectively). Predicted metabolite presence from estimated marginal means showed no significant differences in amino acid breakdown metabolites among sites, as well (Figure 6D, E, and F).

Endogenous protein breakdown metabolite presence varied significantly with collection site in Lake Winnipeg walleye sampled in 2017. Linear models run with hydroxyproline, dimethylarginine, and methionine sulfoxide, potential endogenous protein breakdown biomarkers, were each significant (*F* = 24.12, 6.2, and 6.3, *p* = 1.2 × 10^−8^, 0.0017, and 0.0016, adjusted *R*^2^ = 0.65, 0.29, and 0.29, respectively, Table 5). Moreover, site collected was both a significant independent variable and had a higher effect size than fork length in each model (Table 5). Predicted presence for each candidate metabolite associated with protein breakdown was higher in the northern Dauphin River than the southern Red River (Figure 6G, H, and I). Hydroxyproline and dimethylarginine were also higher in fish from the Dauphin River than at Matheson Island in the channel.

## Discussion

### Morphological Observations

In the current study, we aimed to understand how changing food availability may be affecting walleye and could potentially contribute to large-scale changes in the size and abundance of walleye that have been observed in Lake Winnipeg in recent years. We used the Lake Winnipeg gillnet index data set to examine length-at-age and relative mass to assess trends in the growth of walleye from 2009 to 2018. The decreases in walleye mass appeared to coincide with the near complete collapse of the rainbow smelt population in 2013 as there was a significant drop in relative mass (analogous to body condition) at all sites in the gillnet index between 2014 and 2015. The pattern of overall decrease in mass remained in fish collected in the index until 2018, the last year of data available at the time of analyses. We attempted to control for as many possible confounding variables in the analyses, however the overall trend showed a decrease in relative mass regardless of sex or gillnet mesh size, a pattern that is consistent with previous analyses (Manitoba Government, 2018). Contrary to our predictions, the general trend of decreasing relative mass was not more severe in the north basin (see Figure 5A) where rainbow smelt contributed to a larger portion of the diet of walleye (Sheppard et al., 2015). Given that the decrease in relative mass occurred at all sites sampled, these data suggest ecosystem-wide changes in Lake Winnipeg have contributed to decreases in walleye size over the past decade.

Length-at-age estimates were used as a more stable measurement of changes in growth patterns between 2009 and 2018 because it is less variable than measures of mass that can fluctuate more rapidly (e.g., pre- and post-spawning, post-overwintering). Interestingly, there was a pattern of decreased length in two-year old fish over time, which was most dramatic for fish caught at the Dauphin River site between 2017 and 2018. Reduced food availability or an increase in energetic costs in 2016 and 2017 for young fish at the Dauphin River may have led to this decreased length-at-age. At all sites in the gillnet index, there was also an overall decrease in the length of 6-year old fish from 2016 to 2018. The pattern of decreased length in fish was most prominent in fish collected at the most northern site sampled, Grand Rapids. We believe that this decrease in fish growth in later years (i.e., 2016–2018) of the six-year-old fish may represent a delayed effect from an earlier large-scale change in the ecosystem (e.g., collapse of the rainbow smelt population in 2013). The decrease in the length at age in older fish may have further impacts on walleye abundance in Lake Winnipeg as the length of fish and fecundity is highly correlated (Craig et al., 1995; Wolfert, 1969). Therefore, the population-level impacts of the decrease in the size of walleye in Lake Winnipeg may become a bigger issue in the future.

While the present study is focused on the collapse of the rainbow smelt, additional differences between basins may underlie observed spatial patters. Some of these basin-level differences include higher temperatures, precipitation, river discharge, suspended solids, sulphate, phosphorous, and nitrogen in the south relative to the north basin (Environment Canada, 2011). Phosphorous loading and summer surface temperatures may contribute to algal blooms, which were more prevalent in the south basin (Binding et al., 2018; Environment Canada, 2011). Meanwhile, sodium and chloride are twice as high in the north than in the south, likely because of inflow from the Dauphin River (Environment Canada, 2011). Prey fish populations other than the rainbow smelt also differ spatially, with emerald shiner (*Notropis atherinoides*) and cisco (*Coregonus artedi*) more abundant in the south basin (between 2002-2008) as well as in the diets of south basin walleye (in 2010 and 2011) (Lumb et al., 2012; Sheppard et al., 2015). When these differences between basins are considered in conjunction with data showing that rainbow smelt did not historically make up the entirety of walleye diets (Sheppard et al., 2015), it becomes clear that the rainbow smelt collapse is one of many potential factors that have affected walleye growth. Nevertheless, the disparity in growth rate (length-at-age) between basins, the higher abundance of rainbow smelt in the north basin, and its prevalence in north basin walleye diets in 2010 and 2011 (Sheppard et al., 2015) despite high walleye connectivity between basins (Backhouse-James and Docker, 2011; Thorstensen et al., 2020) suggests the rainbow smelt collapse had a large effect on the Lake Winnipeg walleye fishery.

### Metabolites

We observed from a preliminary analysis of a large-scale targeted metabolomics study (with 163 unique metabolites) that essential amino acids varied in whole blood from walleye caught from different regions of Lake Winnipeg (Wiens, Jeffrey, Treberg unpublished data), which provided the impetus to pursue the nutritional biomarkers strategy that we employed in the current study. We focused on three essential amino acids (methionine, tryptophan and lysine), that were each elevated in the Dauphin River fish relative to the more southern Red River fish (Figure 6). Whole blood was necessary because it was not possible to separate plasma or serum in the field, and we therefore cannot distinguish between differences at the cellular level (mainly red blood cells), or extracellular component of blood. Despite this limitation, because blood acts as a connection between all organs and tissues, we are nevertheless confident that the more north basin walleye had elevated essential amino acids in circulation.

Interpretation of changes in circulating levels of amino acids is complicated by the dynamic nature of amino acid levels in the blood. Most fish nutritional studies focus on the plasma (extracellular component) of blood, and show that essential amino acids may be high due to two mutually exclusive reasons. During periods of high feeding success, the removal of circulating amino acids may not be sufficient to prevent the amino acid levels from elevating in the circulation. In other words, high amino acid levels can reflect high food intake. Alternatively, if the animal must rely on increased protein breakdown due to insufficient feeding, then circulating levels of essential amino acids may also increase (Blasco et al., 1991; Schuhmacher et al., 1995; Costas et al., 2011).

That north basin fish in 2017 displayed lower length-at-age, or less growth, indicates the northern fish may have had lower feeding success in that year compared to the faster growing south basin walleye (Figure 4). Even though the growth estimates support the idea that higher amino acid levels do not reflect greater feeding success for 2017, more information is needed to provide sufficient context for interpreting plasma amino acid levels. Our metabolite screening yielded results for metabolites specific to the degradation pathways each of the three essential amino acids we found to be elevated in the blood of Northern basin fish (Figure 6D, E, and F). Animals use each specific amino acid for energy metabolism by committing each amino acid to its specific degradation pathway. Elevated levels in the metabolites of amino acid degradation therefore imply increased oxidation of each amino acid. Thus, we use these degradation metabolites (dimethyl sulfone, kynurenine, α-aminoadipic acid) as three independent tests of whether or not essential amino acids are being directed towards energy metabolism. In all cases, there was no difference in the metabolites of amino acid oxidation across sampling site. Since most amino acid oxidation takes place outside of the blood (Jürss and Bastrop, 1995; Ballantyne, 2001), one caveat is that blood levels of these metabolites may not sufficiently reflect true whole animal amino acid oxidation. For that reason, the possibility that elevated amino acids reflect a greater reliance on amino acids as energy sources in the Dauphin River fish cannot be outright rejected. However, taken together with the lower growth in the north basin walleye in 2017, we can conclude that it is unlikely the northern walleye have less reliance on protein oxidation than south basin walleye.

Because we focused on amino acids in the blood, and protein turnover is a major source of those circulating amino acids, we needed a means to investigate spatial patterns of protein degradation. To distinguish between amino acids in general and those amino acids that have already been incorporated into proteins, we focused on amino acids that have been modified after they were incorporated into proteins. Three unique protein modifications that arise from three separate routes are focused on: two that come from enzymatic processes (hydroxyproline and dimethylarginine), and one that occurs spontaneously as exposed methionines are oxidized by reactive molecules such as hydrogen peroxide (methionine sulfoxide). While methionine sulfoxide can be formed from free methionine, most methionine from tissues is bound in proteins, so we assume that the bulk of circulating methionine sulfoxide is of proteinaceous origin. Because the means of forming these three post-translational modifications from amino acids represent independent pathways, we had no *a priori* expectation that the levels of all three post-translational modifications would be observed to vary in a consistent fashion across Lake Winnipeg. Nevertheless, all three modifications were higher in fish from the north than the south basin (Figure 6G, H, and I). The signal of protein degradation, based on the consistent presence of post-translationally modified amino acids, could therefore suggest a higher rate of protein turnover in walleye from the north basin of Lake Winnipeg than the south.

If an increased reliance on protein oxidation for energy is *not* driving the elevated amino acids in north basin fish, then the question remains of why there was an increase in protein turnover in those northern fish. While protein metabolism is controlled by many factors, both protein synthesis and degradation rates in fish are known to increase with the level of swimming activity (Houlihan and Laurent, 1987). If the north basin walleye must spend more time swimming to find sufficient nutritional resources, then elevated levels of post-translationally modified amino acids could reflect a greater level of activity for the fish in the north. Relevant to these results, and contrary to what may be intuitive for carnivorous fishes, careful estimations of fuel usage in relation to swimming speed indicates that protein is likely not a major fuel source to supply the increased energy demand of increased swimming in rainbow trout and Nile Tilapia during short-term swimming (Alsop and Wood, 1997; Alsop et al., 1999); although, using the same strategies in Nile Tilapia indicated that prolonged swimming (at ~ 2.7 body lengths/second) over the course of over 48 hours did increase the relative reliance on protein to fuel swimming in unfed fish (Alsop et al., 1999). Since walleye in the wild appear to spend most of their time swimming at speeds below 1.0 body lengths/second (Kelso, 1978), it seems that the extended swimming response in tilapia is not likely applicable to the walleye sampled from the wild in the present study. We therefore suggest that elevated signals of protein turnover in wild walleye may be due to increased swimming activity, which may be undertaken to compensate for decreased food availability.

### Linking Data Sets

The metabolite and gillnet index datasets were consistent in patterns of possible decreased food availability in the north basin in 2017 (when metabolites were measured), based on analyses of spatially varying growth rates in the 2017 gillnet index data and slopes of length-mass relationships in both datasets. Moreover, data set origin (gillnet index or metabolite-measured walleye captured via electrofishing) was not a significant predictor of overall length-mass relationship in a combined linear model, providing additional evidence for no systematic bias in length-mass relationship in one data set or another, even if the walleye used for metabolites were large. Altogether, we argue that the walleye used for assessing metabolite levels are likely representative of large walleye captured in the gillnet index data, and that spatial differences in the slopes of length-mass relationships (≈ body condition) and length-at-age (≈ growth rate) are consistent with spatially varying metabolite presence.

The time scale in response variables should be considered when relating ecology to length-at-age, length-mass relationships, and metabolite levels across data sets. Length-at-age estimates are summaries of one or more prior years of growth for a cohort of walleye, while mass may change from a single feeding or spawning event, thus changing length-mass relationships. In addition, while we do not have information for the response time of amino acids or their metabolites, other metabolites such as blood glucose and lactate increased with handling on a timescale of minutes, indicating that amino acid metabolite presence may also reflect the past several minutes of a fish’s life as opposed to ecological patterns (Chopin et al., 1996; Grutter and Pankhurst, 2000; Meka and McCormick, 2005; Lawrence et al., 2018). However, the consistency in patterns across timescales— stronger signals of protein breakdown, more shallow length-mass relationships, and slower growth in the north basin in 2017—supports the validity of this approach for integrating data from different levels of biological organization. The breadth in timescales analyzed may also be a benefit for integrating information across data sets because each piece of information provides context for other results.

### Conclusions

The data presented show declining growth rates and condition in Lake Winnipeg walleye in recent years, especially those in the north basin. These morphological differences are consistent with both the collapse of the rainbow smelt population and blood metabolites, suggesting increased endogenous protein breakdown in the north basin. With validation and refinement, the metabolites identified in the present study thus have potential for further development into molecular markers possibly useful as indicators of nutritional status for the walleye fishery. Molecular indicators of nutritional status would be valuable tools for resource managers for describing physiological thresholds in nutritional status that are predictive of detrimental effects on the walleye fishery (Connon et al., 2018). In other words, a molecular panel describing nutritional status may support the sustainable management of the Lake Winnipeg fishery.

## Acknowledgments

We thank E. Enders, D. Watkinson, C. Charles, C. Kovachik, D. Leroux, N. Turner, M. Gaudry, S. Glowa, and E. Barker for their role in sampling the walleye used for metabolites. C. Charles and E. de Greef assisted with a map of Lake Winnipeg, and E. de Greef also provided immense support in the process of writing the manuscript. This work was supported by a Fisheries and Oceans Canada Ocean and Freshwater Science Contribution Program Partnership Fund grant awarded to J.R.T., K.M.J. and Darren Gillis, and Natural Sciences and Engineering Research Council of Canada Discovery Grants awarded to K.M.J. (#05479) and J.R.T. (#06052). Work by J.R.T. is also supported by the Canada Research Chairs program (#223744) and the Faculty of Science, University of Manitoba (#319254).

